# Visuomotor learning promotes visually evoked activity in the medial prefrontal cortex

**DOI:** 10.1101/2022.05.31.494126

**Authors:** Andrew J. Peters, Andrada-Maria Marica, Julie M.J. Fabre, Kenneth D. Harris, Matteo Carandini

**Affiliations:** UCL Institute of Ophthalmology, University College London, London, UK; UCL Queen Square Institute of Neurology, University College London, London, UK

## Abstract

The medial prefrontal cortex (mPFC) is necessary for executing many learned associations between stimuli and movement. It is unclear, however, whether activity in the mPFC reflects sensory or motor aspects of sensorimotor associations and whether it evolves gradually during learning. To address these questions, we recorded cortical activity with widefield calcium imaging while mice learned a visuomotor task. The task involved associating a visual stimulus with a forelimb movement. After learning, the mPFC showed stimulus-evoked activity both during task performance and during passive viewing, when the stimulus evoked no action. This stimulus-evoked activity closely tracked behavioral performance across training, exhibiting jumps between training days. Electrophysiological recordings localized this activity to the secondary motor and anterior cingulate cortex. We conclude that learning a visuomotor task promotes a route for visual information to reach the prefrontal cortex, which develops responses to the relevant visual stimuli even outside the context of the task.

## INTRODUCTION

The medial prefrontal cortex (mPFC), consisting of the secondary motor, anterior cingulate, prelimbic, and infralimbic cortex (Le Merre et al., 2021), is a nexus of sensory and motor information (Harris et al., 2019) and has been suggested to orchestrate activity across the brain (Allen et al., 2017; Makino et al., 2017). Accordingly, inactivation of the mPFC impairs sensory-guided movements, indicating that it is causally involved in transforming stimuli into actions (Pinto and Dan, 2015; Siniscalchi et al., 2016; Zatka-Haas et al., 2021). The mPFC is therefore thought to be critical for learning arbitrary associations between stimuli and movements (Crochet et al., 2019; Le Merre et al., 2021; Murray et al., 2000).

Supporting the role of the mPFC in learning sensorimotor associations, the mPFC exhibits sensory responses specifically to stimuli that are behaviorally relevant (Bichot et al., 1996; Wal et al., 2021). These stimuli can be of any modality including visual (Orsolic et al., 2021; Peters et al., 2021; Reinert et al., 2021), auditory (Moorman and Aston-Jones, 2015; Pinto and Dan, 2015), somatosensory (Le Merre et al., 2018), and even multisensory (Coen et al., 2021).

While it is clear that activity in the mPFC changes after learning (Mulder et al., 2003; Orsolic et al., 2021; Otis et al., 2017; Peters et al., 2021), it is less clear what happens during learning. It has been proposed that learning proceeds apace with an increase in sensory-evoked mPFC local field potentials (Le Merre et al., 2018) and that the mPFC exerts a gradually increasing influence over other cortical areas (Makino et al., 2017).

Moreover, the exact location of mPFC stimulus responses is unclear, as the mPFC spans multiple regions (Le Merre et al., 2021). For example, while frontal regions along the cortical midline can respond to visual stimuli even without training (Mohajerani et al., 2013; Murakami et al., 2015; Sreenivasan et al., 2016a), it may be that learning drives responses to trained visual responses in more anterior regions (Orsolic et al., 2021; Peters et al., 2021; Reinert et al., 2021).

Here we address these questions by performing longitudinal widefield calcium imaging in the mouse dorsal cortex throughout learning of a visuomotor association. We find that stimulus responses arise selectively to the trained stimulus after learning, specifically in the dorsomedial prefrontal cortex (dmPFC, defined to mean anteromedial secondary motor and underlying anterior cingulate cortex). Those responses are selective to the learned stimuli, occur in passive viewing as well as task performance, arise from the first day of learning, and increase with subsequent training, displaying an intriguing series of jumps across training days. Together, our results demonstrate visuomotor association learning is closely tracked by the emergence of task stimulus responses in the dmPFC.

## RESULTS

We trained mice in an operant task requiring a single stimulus-movement association (**Figure 1A**). Mice were surrounded by screens on their left, front, and right, and rested their forepaws on a steering wheel. Visual grating stimuli appeared on the right-hand screen and could be moved leftwards by turning the steering wheel counterclockwise to obtain a sucrose water reward. Turning the wheel clockwise moved the stimulus rightward off the screen and triggered a burst of white noise. In a rewarded trial, the stimulus was fixed in place on the center screen for one second while the mouse consumed the sucrose. This task is derived from a 2-alternative choice task that we have developed previously (Burgess et al., 2017; International Brain Laboratory et al., 2021) but it is simpler, because the stimulus is always high-contrast and always appears on the right side. The interval between stimulus presentations consisted of a fixed inter-trial interval, followed by a quiescence period, which restarted if the wheel was turned. In each trial, the delay parameters – the inter-trial interval and quiescence period – were chosen randomly, so that the stimulus appeared at an unpredictable time, and only while the mouse was not moving the wheel.

**Figure 1.**
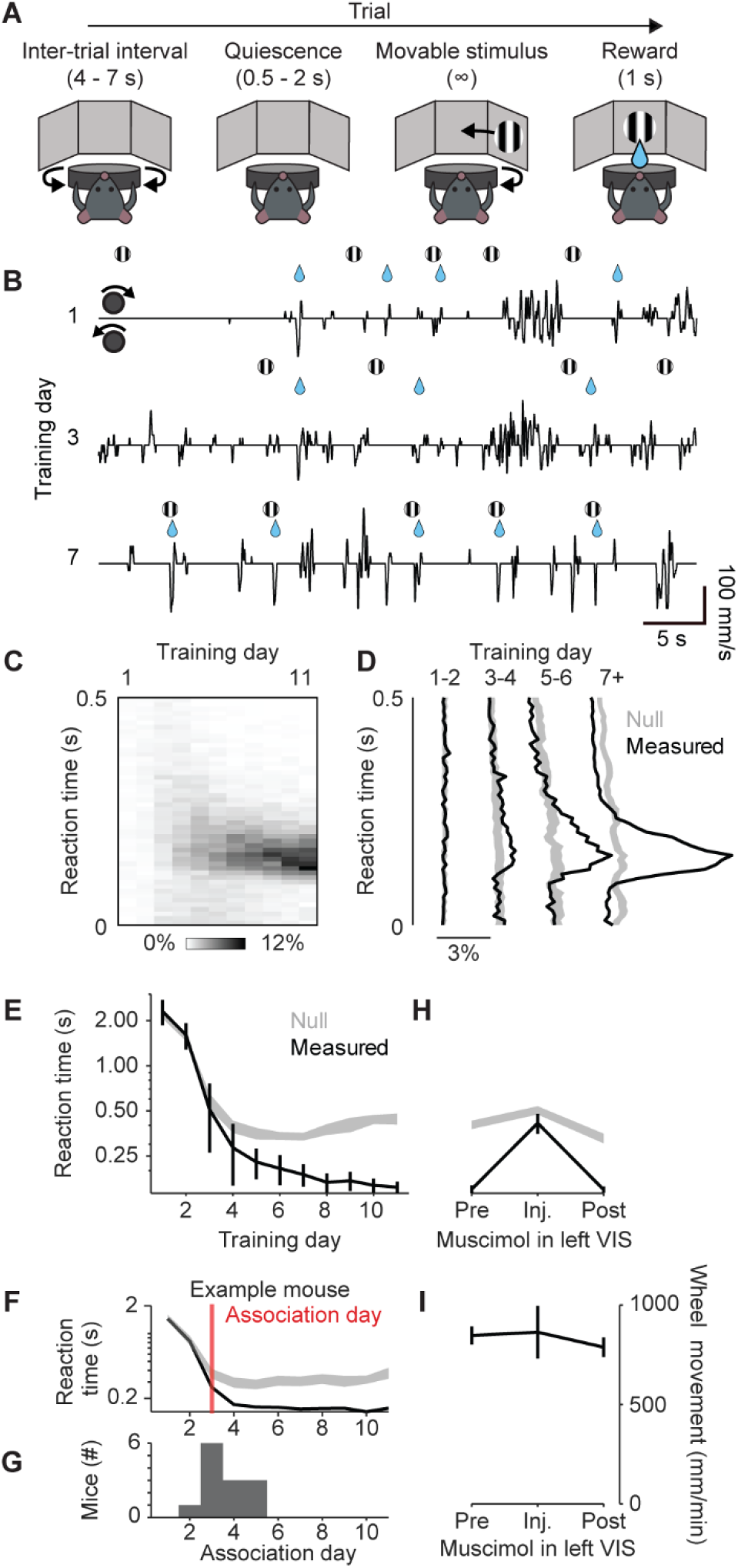
Visuomotor association task. (A) Task trial structure. (B) Example segments of task events and wheel movement from one mouse across three days. (C) Stacked histograms of reaction times as a function of training day, averaged across all mice (n = 13 mice). (D) Histograms of reaction times at different learning points *(black),* compared to prediction from chance *(gray).* Curves are mean across mice; shadings, 95% confidence intervals of the null distribution (n = 13 mice) (E) Median reaction times for each training day (*black*) compared to prediction from chance *(gray).* Curves and error bars show median ± m.a.d. across mice (n = 13 mice), shading shows 95% confidence intervals from the null distribution. (F) Median reaction times as in (E) with “association day” for one example mouse. (G) Histogram of association day across mice. (H) Effect of V1 muscimol on reaction times in trained mice, plotted as in E. Inactivating V1 reversibly increases median reaction times (one-way ANOVA, p = 3.3 x 10^-1^). (I) Effect of muscimol on total wheel movement. Curves and error bars show mean ± s.e. across mice (n = 5 mice). Wheel movement is not different across groups (one-way ANOVA, p = 0.81).

Mice learned to robustly associate the right-hand visual stimulus with a counterclockwise wheel turn in less than one week. They learned to move the wheel counterclockwise on 81 ± 14% (s.d., n = 13) of trials on the very first day, but the timing of these turns was unrelated to visual stimulus onsets (**Figure 1B**, *days 1 and 3*). After further training, they began to reliably turn the wheel 100-200 ms after stimulus onset (**Figure 1B**, *day 7),* becoming more locked to visual stimuli over progressive days (**Figure 1C**).

To statistically analyze this sharpening of reaction times, we developed a conditional randomization method. We compared the actual reaction times to a null distribution that was derived from the same wheel turn trajectories, but with randomized stimulus onset times (obtained by randomly resampling from the set of delay parameters that would have led to the same detected movement time as actually observed; STAR Methods). Under the null hypothesis that actions are unrelated to the visual stimuli, the reaction times would be a random sample from this null distribution.

This method showed that an initial decrease in reaction times observed over the first few training days was seen also in the null distribution, as it simply reflected an increased propensity to turn the wheel at times unrelated to the visual stimulus (**Figure 1D-E**). In later days, however, a peak emerged at 100-200 ms post-stimulus time peak, which was not also seen in the null distribution, indicating time-locking to the visual stimulus. Using this approach, we could establish the “association day” for each mouse: the first day when its reaction times diverged from the null distribution (**Figure 1 F**). Most often, this was the 3^rd^ day of training (**Figure 1G, Figure S1**).

Even though this task involves high-contrast visual stimuli, which could in principle be processed subcortically (Glickfeld et al., 2013), the visual performance of the mice depended critically on the visual cortex. In a subset of mice (n = 5), we injected muscimol into the left visual cortex, thus eliminating visual responses in the left hemisphere (**Figure S2A**). This inactivation increased reaction times to chance levels (**Figure 1H**, **S2B**), but did not affect total wheel movement (**Figure 1I**). Thus, mice were equally engaged in the task with or without visual cortical activity, but when the visual cortex was inactivated, they were greatly impaired at responding specifically to the visual stimulus.

Visually evoked activity emerged in the mPFC with task learning. We performed widefield imaging on all training days, in mice expressing GCaMP6s in excitatory neurons (Wekselblatt et al., 2016). Fluorescence in these mice can be seen in the entire dorsal cortex, and after deconvolution this signal correlates well with the spiking of neurons in deep layers (Peters et al., 2021). To examine how cortical activity changes with learning, we averaged responses in all days before each mouse’s association day to represent the novice stage, and all days after the association day to represent the trained stage. In both stages, we observed activity in the visual cortex after stimulus onset, and the forelimb somatomotor, retrosplenial, and medial prefrontal cortex (mPFC) at movement onset (**Figure 2A**). While the broad patterns were similar in novice and trained stages, a key difference was seen in the left hemisphere mPFC, which exhibited increased stimulus-evoked activity in trained mice (**Figure 2A,** *bottom left).* The predominantly unilateral nature of this response matched that of visual cortex, suggesting a response to the right-hand visual stimulus.

**Figure 2.**
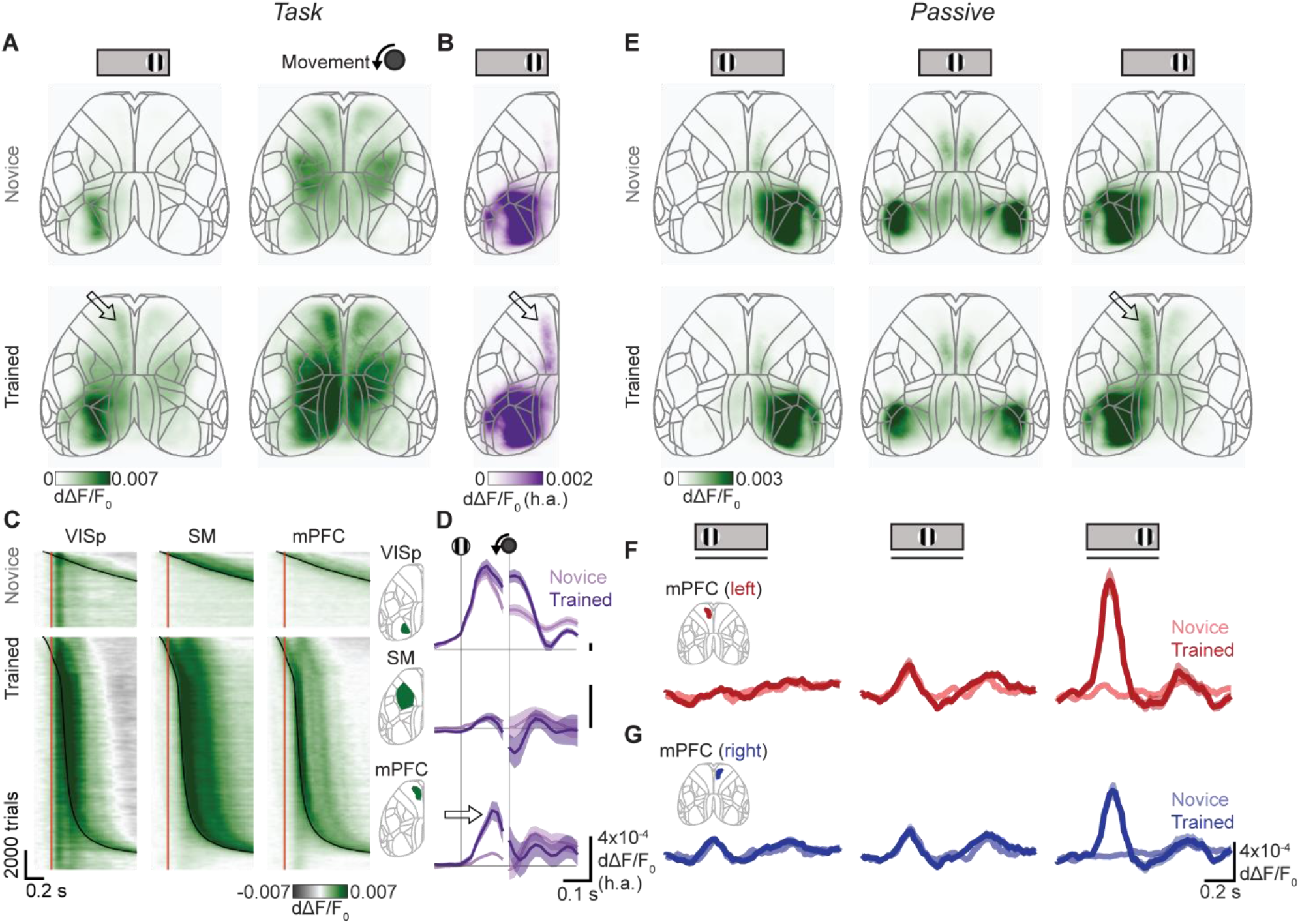
mPFC develops stimulus-evoked responses after learning. (A) Mean fluorescence 100 ms after stimulus onset (left) and 0 ms after movement onset (right), for novice (top) or trained (bottom) mice. Values are deconvolved fluorescence relative to baseline (dΔF/F0). Arrow: left hemisphere mPFC activity after learning. (B) Average stimulus response across mice, computed as the hemispheric asymmetry (h.a.) of the maximum fluorescence 0-200 ms after stimulus onset, to remove bilateral movement-related activity. Arrow: left hemisphere mPFC activity after learning. (C) Fluorescence in left primary visual cortex (VISp), limb somatomotor cortex (SM), and medial prefrontal cortex (mPFC) for all trials and mice, aligned to stimulus onset and sorted by training stage (top, novice; bottom, trained) and reaction time. Red lines, stimulus onset; black curves, movement onset. (D) Hemispheric asymmetry of fluorescence for novice (light purple) or trained (dark purple) mice in the same three regions of interest. Curves and shading show mean ± s.e. across mice, positive means more activity in the left hemisphere. Arrow: maximum fluorescence 0-200ms after stimulus onset increases only in the left mPFC after learning (one-way ANOVA, VISp p = 0.33, SM p = 0.72, mPFC p = 2.7 x 10^-4^). (E) Maximum fluorescence 0-200 ms after stimulus onset in novice (top) and trained (bottom) mice, to visual stimuli on the left, center and right (columns). (F) Fluorescence during passive stimulus viewing in left hemisphere mPFC in novice (light red) and trained (dark red) mice. Curves and shading show mean ± s.e. across mice. Line under stimulus icon indicates when the stimulus is on the screen. The left mPFC has an increased response to contralateral (right-hand) stimuli after learning (three-way ANOVA on time, learning stage, and stimulus, learning stage effect p = 2.8 x 10-^26^). (G) As in (F), for the right hemisphere mPFC. The right mPFC has a small response to contralateral (left-hand) stimuli which does not change with learning (two-way ANOVA on hemisphere and learning stage, hemisphere effect p = 1.5e-5, learning stage effect p = 0.37; responses measured as maximum 0-200 ms after stimulus onset). The right mPFC also responds to ipsilateral (right-hand) stimuli but only after learning (three-way ANOVA on time, stage, and stimulus, stage effect p = 3.5 x 10^-19^).

This visual activity in the mPFC was followed by motor activity (**Figure 2A**, *right),* which acts as a confound because (by definition) trained mice performed more movements in association with the stimulus. To minimize this confound we exploited the fact that the visual response is predominantly unilateral whereas movement responses and spontaneous fluctuations are bilateral (Shimaoka et al., 2019). We thus estimated the hemispheric asymmetry of the response: we subtracted right hemisphere fluorescence from left hemisphere fluorescence, after weighting by a constant fit by linear regression during movement with no stimuli (STAR Methods). The resulting maps indicated strong asymmetrical stimulus responses in visual cortex, which were expected and unchanged by learning, and revealed asymmetric stimulus responses that appeared in the mPFC only after learning (**Figure 2B**, *arrow).*

The increase in visually evoked activity after learning was unique to the mPFC and was not obviously related to task movements. In both the novice and learned stages, the primary visual cortex (VISp) exhibited stimulus-aligned activity (**Figure 2C**, *left*) while the somatomotor cortex (SM) exhibited movement-aligned activity (**Figure 2C**, *middle).* The mPFC, on the other hand, exhibited only movement-aligned activity before learning but showed additional stimulus-aligned activity after learning (**Figure 2C**, *right*). As before, we could isolate stimulus-evoked activity through a weighted hemisphere difference. Stimuli evoked a consistent response in the left visual cortex regardless of learning (**Figure 2D**, *top)* and no response in the somatomotor cortex (**Figure 2D**, *middle*), while the mPFC gained a new stimulus-evoked response after learning (**Figure 2D**, *bottom, arrow).*

Learning a visuomotor association therefore appeared to drive the development of stimulus-evoked activity in the mPFC. During the task, however, this stimulus-evoked activity overlaps with movement-related activity, and this overlap makes it difficult to precisely distinguish the two, and more generally raises the question of whether the two forms of activity are related to each other. To address these issues, we next analyzed activity measured during passive stimulus viewing, when the mouse viewed similar stimuli but did not perform any action.

After learning, the mPFC exhibited visually evoked activity even during passive stimulus viewing, and this activity was specific to the trained stimulus. We presented stimuli on the left, center, and right screens to passive mice, for three days prior to training, and after every training session. To avoid contamination with movement responses, all trials with wheel movement were removed from analysis. Activity in the visual cortex appeared in the hemisphere contralateral to left or right stimuli, or bilaterally for central stimuli, and was present throughout training but became more sustained after learning (**Figure 2E, Figure S3**). A region in the posterior secondary motor cortex was also consistently responsive across novice and trained stages, as previously observed (Mohajerani et al., 2013; Murakami et al., 2015; Sreenivasan et al., 2016b) (**Figure 2E**). The mPFC responded weakly to contralateral stimuli in novice mice, but developed a strong response after learning, specifically to the right-hand stimulus which animals had learned to detect during the task (**Figure 2E-G**, *right column).* Indeed, the left mPFC responded to neither the left-hand stimulus (never viewed during the task) nor the central stimulus (viewed during reward in the task) (**Figure 2E-F**, *left and center columns*). The mPFC response to right-hand stimuli was present in both hemispheres, although it was much larger in the left hemisphere contralateral to the stimulus (**Figure 2F-G**, *right column).* These mPFC stimulus responses depended on an intact visual cortex, as they were eliminated when visual cortex was inactivated with muscimol (**Figure S2C-E**). Thus, learning-induced stimulus responses in the mPFC were not specific to task engagement, and were downstream of responses in the visual cortex.

The mPFC responses evoked by visual stimuli in passive conditions could not be explained by subtle movements elicited by task stimuli. Passive stimulus presentation did not evoke overt limb movements, although video analysis of the face and eye showed that mice could exhibit subtle behavioral changes after learning, including a pupil dilation and slight whisker twitch (**Figure S4A-D**). We took advantage of substantial trial-by-trial variability to bin trials according to whisker movement, allowing us to find trials with similar amounts of whisker movement in both the novice and trained stages (**Figure S4E**). These variable movements were accompanied by consistently low mPFC fluorescence in novice days and high mPFC fluorescence after learning (**Figure S4F-G**) indicating that visually evoked mPFC responses were increased beyond what could be accounted for through behavior.

Visually evoked widefield responses in the mPFC closely tracked behavioral performance across days. Because different mice learned the task at different rates, we examined the timecourse of activity changes relative to the association day, where performance first exceeds chance. Stimulus-evoked mPFC activity increased substantially on the association day during both task performance and passive stimulus viewing after that training session (**Figure 3A-B**). Remarkably, neither reaction times nor stimulus-evoked mPFC activity increased over the course of a day’s training, instead jumping between days to attain values outside of those from each previous day (**Figure 3CD**). After the association day, reaction time continued to decrease, accompanied by increases in mPFC stimulus responses, with both reaching a plateau approximately five days after the association day (**Figure 3C-D**). Together, these results show that stimulus-evoked activity in the mPFC mirrored behavioral learning, with both jumping in a stepwise fashion between training days.

**Figure 3.**
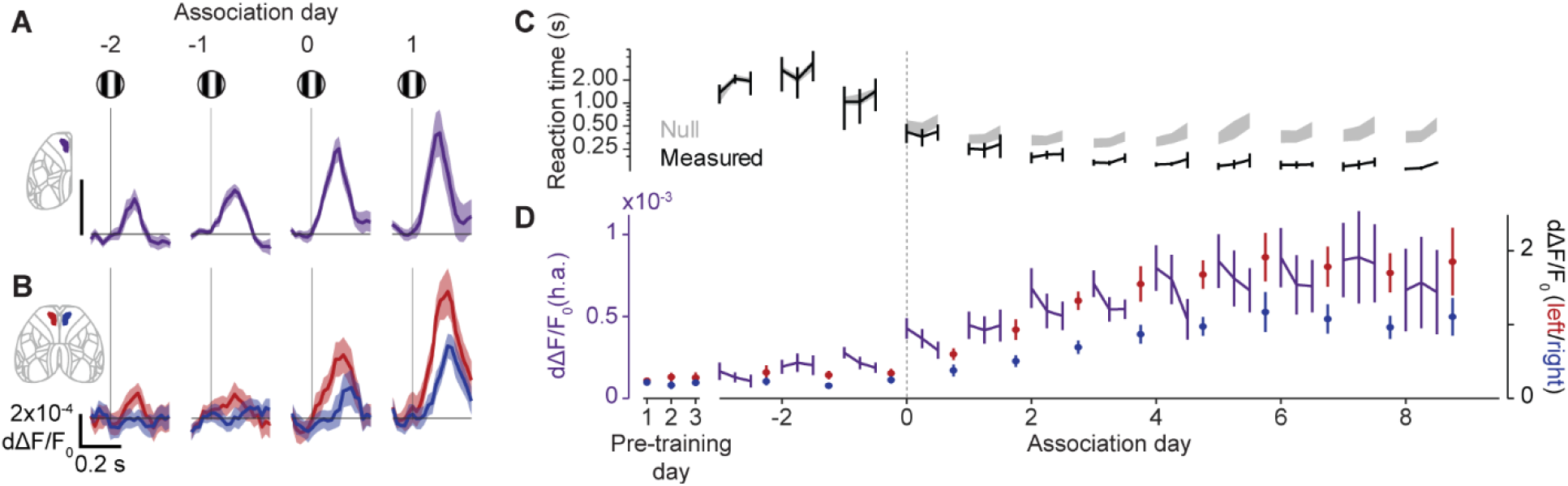
mPFC stimulus-evoked responses track behavioral performance in jumps across days. (A) Fluorescence in the mPFC as hemispheric asymmetry during task performance, aligned to stimulus onset and plotted relative to association day. Positive means more fluorescence in the left hemisphere, day 0 is the association day. Curves and shading show mean ± s.e.m. across mice (n = 13 mice). Maximum fluorescence 0-200 ms after stimulus onset increases specifically on the association day (signed-rank test, day −2 vs −1 p = 0.23, day −1 vs 0 p = 6.1 x 10^-3^.) (B) Fluorescence in the left (red) and right (blue) mPFC during passive viewing of right-hand stimuli. Curves and shading show mean ± s.e.m. across mice (n = 13 mice). Maximum fluorescence 0-200 ms after stimulus onset increases specifically on the learned day (signed-rank test, day −2 vs −1 p = 0.62, day −1 vs 0 p = 2.4 x 10^-3^). (C) Median reaction times as function of day relative to association day. For each day, the three connected points represent three equal thirds of each day. Curves and error bars show median ± m.a.d. across mice (n = 13 mice), shading shows 95% confidence intervals from the null distribution. (D) Maximum fluorescence 0-200 ms after stimulus onset in the mPFC relative to association day, during the task as hemispheric asymmetry (h.a., purple) and during passive viewing of right-hand stimuli in the left (blue) and right (red) hemisphere. Task fluorescence is split into thirds of trials for each day, and passive fluorescence is included for three days prior to training. Curves and error bars show mean ± s.e.m. across mice (n = 13 mice).

After learning, mPFC neurons encoding the stimulus were found mainly in the secondary motor and anterior cingulate cortex. To determine which regions within the mPFC contained cells with stimulus-evoked activity, we used acute electrophysiological recordings in trained mice (10 recordings across 5 mice, all of whom had previously undergone widefield imaging). We targeted Neuropixels probes to the area exhibiting stimulus responses after learning based on widefield fluorescence, and frequently observed neurons that responded to both stimuli and movement, as quantified through passive stimulus presentation (without movement) and delay-period movements during the task (without visual stimuli) (**Figure 4A**). We histologically verified that the probes passed through the secondary motor (MOs), anterior cingulate (ACA), prelimbic (PL), and infralimbic (ILA) cortex (**Figure 4B**). Multiunit signals showed visual responses selectively to the trained right-hand stimulus, in a decreasing dorsal to ventral gradient, with the largest responses in secondary motor and anterior cingulate cortex (**Figure 4C**). The mPFC activity we observed with widefield imaging thus originates primarily from the dorsomedial PFC (dmPFC): the secondary motor and the anterior cingulate cortex.

**Figure 4.**
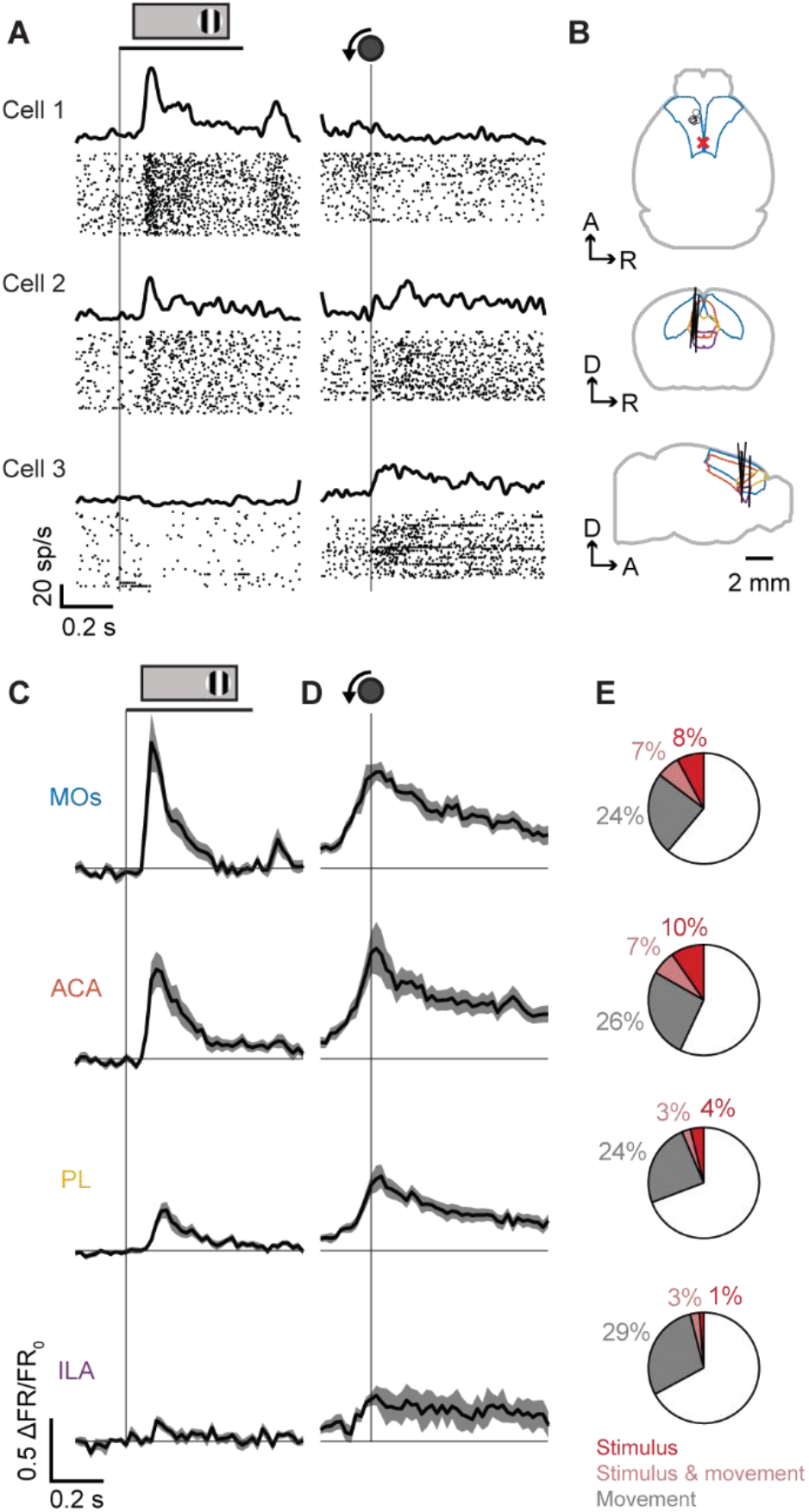
mPFC neurons of trained mice show stimulus- and movement-evoked responses in secondary motor and anterior cingulate cortex. (A) Raster plots for example cells aligned to passive right-hand stimulus presentation (without movement; left column) and movement onset during task delay periods (without visual stimuli; right column). (B) Neuropixels probe locations (black lines) and outlines of secondary motor (blue), anterior cingulate (orange), prelimbic (yellow), and infralimbic (purple) cortex. Red x represents bregma. Each black line is from one mouse (n = 5 mice). (C) Multiunit responses to passive viewing of right-hand stimuli for four frontal areas, including secondary motor (MOs), anterior cingulate (ACA), prelimbic (PL), and infralimbic (ILA) cortex. Curves and shading show mean ± s.e.m. across recordings. Line under stimulus icon indicates when the stimulus is on the screen. (D) Multiunit responses to movement onset during delay periods of the task (without stimuli), from the same recording days and areas in (C). Curves and shading show mean ± s.e.m. across recordings. (E) Fraction of neurons responding positively or negatively to right-hand stimuli during passive viewing (red; p < 0.01, shuffle method), to movement onset during task delay periods with no stimulus (gray), and to both (pale red). Values represent mean across recordings (n = 10 recordings). Stimulus-responsive cells are more likely to be movement-responsive than expected from chance in the dmPFC (shuffling movement-responsive classification within area, p < 1 x 10-^4^).

The visual responses of dmPFC were carried by few neurons, and these neurons were often also responsive to movement. Multiunit activity in the dmPFC was evoked by delay-period movements in addition to stimuli, indicating a mixing of sensory and motor responses within this region (**Figure 4D**). To determine whether responses to stimuli and movements were present in the same cells, we quantified responsiveness for high-quality single units. Stimulus responses were rare across units in the dmPFC (16 ± 10% s.d., n = 10 recordings), while movement responses were more common (32 ± 12% s.d., n = 10 recordings). Stimulus-responsive units, however, were more likely to be responsive to movement than expected from chance (**Figure 4E**). These results indicate that the stimulus-evoked response observed in the mPFC arises from a subset of neurons with multi-functional responses.

## DISCUSSION

Our results show that the mPFC becomes responsive to a visual stimulus concurrently with learning a visuomotor association, and that both performance and visual responses in mPFC increase between training sessions, but not within sessions. These findings suggest that stimulus information becomes routed to the mPFC through learning, which in turn may drive execution of the associated movement.

Activity within the mPFC is necessary for mice to execute this type of association, as demonstrated by optogenetic inactivation (Coen et al., 2021; Zatka-Haas et al., 2021). The timing of inactivation effects suggest that stimulus-related activity, rather than movement-related activity, is particularly critical for successful performance (Zatka-Haas et al., 2021). This supports the notion that the behavioral relevance of a stimulus is tied to the propagation of sensory information from the visual cortex to the mPFC.

Our findings reinforce previous work indicating that learning causes the mPFC to respond to stimuli of multiple modalities (Le Merre et al., 2018; Orsolic et al., 2021; Otis et al., 2017; Reinert et al., 2021; Siniscalchi et al., 2016). Furthermore, inactivation of the mPFC in an audiovisual task impairs behavior relating to both auditory and visual stimuli (Coen et al., 2021), indicating a convergence of relevant stimulus information. Multiple areas within the mPFC have been previously associated with stimulus “value” as its reward-predicting capacity, including in MOs (Sul et al., 2011), ACA (Kennerley et al., 2011), and PL (Lak et al., 2020). Nevertheless, we did not observe mPFC responses to central stimuli, even those these stimuli were present whenever reward was delivered to the mice. Our data therefore broadly fit of the idea that the mPFC provides context for movement (Barthas and Kwan, 2017; Heilbronner and Hayden, 2016; Rushworth et al., 2011), and show that such context appears on the first day that visuomotor responses are behaviorally evident. These responses are specific to the learned stimuli, but are not specific to the task context, also occurring in passive stimulus presentation. Thus, these results suggest that plasticity either upstream of or in mPFC has led to representation of task stimuli specifically.

By following activity across learning, we found across-day rather than within-day changes in both behavior and activity, suggesting that learning involves plasticity after training. While some types of association can be developed within a session (Komiyama et al., 2010), it is common to observe across-day changes in neural activity (Costa et al., 2004). These across-day changes have been suggested to result from spine growth and/or activity replay after each day’s training (Peyrache et al., 2009; Yang et al., 2014), which might underlie the jumps in mPFC activity we observe. In our task, mice begin turning the wheel on the first day, but at times unrelated to the visual stimulus, indicating that animals learn to produce this movement even before a sensorimotor transformation between stimulus and movement develops. These untimed movements occurring early in training are also accompanied by mPFC activity. We hypothesize that after early training, mPFC activity is sufficient to drive movements, and that synaptic plasticity occurring in between later training sessions routes visual activity to the mPFC, causing the stimulus to trigger the movement, reflected in an across-day jump in both visually driven behavior and visually driven mPFC activity.

The mPFC is likely part of a larger network that together drives the movement in response to the stimulus. We found that sensory and motor responses overlapped in mPFC regions and even within neurons, in line with previous findings in the prelimbic area PL (Pinto and Dan, 2015). These responses may serve multiple functions downstream, for example with mPFC sensory-aligned activity influencing perception across modalities including visual (Huda et al., 2020; Zhang et al., 2014), auditory (Schneider et al., 2018), and somatosensory (Manita et al., 2015), and mPFC movement-aligned activity coordinating motor commands within the motor cortex (Allen et al., 2017; Makino et al., 2017), superior colliculus (Huda et al., 2020), and striatum (Otis et al., 2017; Peters et al., 2021).

The exact region within the dmPFC (including the secondary motor and anterior cingulate cortex) that developed stimulus responses after learning is separable from many other frontal regions of interest. The learning-related stimulus responses we observed were spread from approximately 1-2.5 mm anterior and 0.2-1 mm lateral to bregma, which is on the medial edge of frontal responses often associated with delay-period activity and higher-order task and movement variables (Svoboda and Li, 2018). Progressing from lateral to medial, this is distinct from the orofacial anterior-lateral motor region (ALM) (Komiyama et al., 2010), likely distinct from the rostral forelimb area (RFA) (Tennant et al., 2011), vibrissal motor cortex (vM1) (Ferezou et al., 2007), and medial motor area (MM) (Chen et al., 2017), but possibly overlapping with the frontal orienting fields (FOF) found in rats (Erlich et al., 2011). The location where we saw learning-related dmPFC responses is also separable from a more posterior medial frontal region that exhibited responses to visual stimuli in novice mice (Mohajerani et al., 2013; Murakami et al., 2015; Sreenivasan et al., 2016a), demonstrating two functionally distinct visually responsive medial frontal regions. It seems unlikely that the precise area in which we found stimulus responses is related to the forelimb being the associated effector, given a previous demonstration that this region is involved in a licking visuomotor task (Goard et al., 2016). This region may therefore represent a specific part of the dmPFC with a unique relationship to learning.

Together, these results demonstrate that learning a visuomotor association is strongly coupled with the emergence of stimulus-evoked activity in the dmPFC, and suggests that the dmPFC is a critical node for tying sensory information to movement.

## STAR METHODS

### Key resources table

| REAGENT or RESOURCE | SOURCE | IDENTIFIER |
| --- | --- | --- |
| Experimental models: Organisms/strains |  |  |
| Mice tetO-GCaMP6s | The Jackson Laboratory | Jax #024742, RRID:IMSR_JAX:024742 |
| Mice CaMK2a-tTa | The Jackson Laboratory | Jax #007004, RRID:IMSR_JAX:007004 |
| Software and algorithms |  |  |
| MATLAB > R2018a | MathWorks | https://www.mathworks.com |
| Rigbox | Bhagat et al., 2020 | https://github.com/cortex-lab/Rigbox |
| Open Ephys GUI | Open Ephys | https://open-ephys.org/gui |
| Kilosort 2.0 | Stringer et al., 2019 | https://github.com/MouseLand/Kilosort/releases/tag/v2.0 |
| Phy | Rossant et al., 2016 | https://github.com/cortex-lab/phy |
| DeepLabCut | Mathis et al., 2018 | https://github.com/DeepLabCut/DeepLabCut |
| AP_histology | Peters et al., 2021 | https://github.com/petersaj/AP_histology |
| Bombcell | This paper | https://github.com/Julie-Fabre/bombcell |
| Chemicals, peptides, and recombinant proteins |  |  |
| Muscimol | Sigma Aldrich | CAS number: 2763-96-4 |
| Dil | ThermoFisher Scientific | Catalog number: V22885 |
| Other |  |  |
| Norland fast-curing optical adhesive | Norland Products | NOA 81 |

### Lead contact and materials availability

Further information and requests for resources should be directed to and will be fulfilled by the Lead Contact, Andrew Peters.

### Experimental model and subj ect details

All experiments were conducted according to the UK Animals (Scientific Procedures) Act 1986 under personal and project licenses issued by the Home Office. Mice were adult (6 weeks or older) male and female transgenic mice (tetO-G6s;Camk2a-tTa, from Ref. (Wekselblatt et al., 2016)).

### Method details

#### Surgery

Surgery for widefield imaging involved affixing a headplate and plastic well to the skull and applying an optically transparent adhesive to the skull. Mice were anesthetized with isoflurane, injected subcutaneously with Carprieve, and placed in a stereotaxic apparatus on a heat pad. The head was then shaved, the scalp cleaned with iodine and alcohol, and the scalp and periosteum were removed to expose the skull. The cut skin was sealed with VetBond (World Precision Instruments), and a custom steel headplate was fixed to the interparietal bone with dental cement (Super-Bond C&B). A plastic 3D-printed U-shaped well was then cemented to enclose the edges of the exposed skull. A layer of VetBond was applied to the skull followed by two layers of UV-curing optical adhesive (NOA81, Norland Products). Carprieve was added to the drinking water for 3 days after surgery.

For muscimol injections, the retinotopically peripheral zone of V1 in the left hemisphere (contralateral to the trained stimulus) was targeted using widefield retinotopic mapping relative to vasculature. Mice were anesthetized with isoflurane, injected subcutaneously with Carprieve, and head-fixed using the previously implanted headplate. The optical adhesive was drilled away over the targeted location and a small craniotomy was drilled. Muscimol (Sigma, 5 mM in ACSF) was injected through a sharpened ~40 μm borosilicate capillary with a pneumatic injector (Nanoject, Drummond Scientific) in two boluses of 70nl at 300 and 700 μm from the cortical surface. The craniotomy was then filled with Kwik-Sil (World Precision Instruments), a thin layer of clear dental cement was applied, and overlying optical adhesive was replaced. Recordings were performed at least 1.5 hours after muscimol injections.

For electrophysiological recordings, the mPFC in the left hemisphere was targeted using widefield responses to right-hand stimuli relative to vasculature (centered at approximately 1.7 mm AP and 0.7 mm ML relative to bregma, Figure 4B). Mice were anesthetized with isoflurane, injected subcutaneously with Carprieve, and head-fixed using the previously implanted headplate. The optical adhesive was drilled away over the targeted location and a small craniotomy was drilled. The craniotomy was covered with Kwik-Cast (World Precision Instruments), and recordings were performed at least 1.5 hours after surgery.

#### Visuomotor operant task

Mice performed an operant task where a visual stimulus on the right-hand side could be moved with a wheel into the center to receive a sucrose reward. Mice were water restricted and typically received their required water amounts during the task, or were otherwise supplemented to a minimum relative to the body weight of each mouse. The task was a variant of one described previously (Burgess et al., 2017) and programmed in Signals, part of the Rigbox MATLAB package (Bhagat et al., 2020).

Visual stimuli consisted of square gratings with 100% contrast, 1/15 cycles per degree, and randomized phase on each trial, within a circular gaussian window of σ = 20° which effectively covered an entire screen. At the start of each trial, the visual stimulus appeared on the right-hand screen and was positionally yoked to wheel movements, for example with counterclockwise (leftward) movements of the wheel bringing the visual stimulus leftward towards the center. If the stimulus was brought to the center a 6 μL sucrose reward was delivered, and if the stimulus was instead moved rightward off the screen a low burst of white noise was played through speakers under the screens. Mice quickly learned to move the wheel leftward instead of rightward, with 81 ± 14% of rewarded trials on day 1 and 96 ± 3% across-session average (mean ± s.d. across 13 mice).

Between the response of one trial and the appearance of the visual stimulus on the next trial, two delay parameters were independently and randomly selected for each trial. The first, an inter-trial interval (ITI), was a fixed time from the outcome of the previous trial. The second, an enforced quiescence period, was a timer that began after the ITI and would reset with any wheel movement. Delay timings were shorter at the start of training and were lengthened once mice first obtained their full daily water amount in the task, which was usually on the first or second day of training. Delay timings were selected from a range in 100 ms increments, with initial ranges being 1-3 s for ITIs and 0.5-1 s for quiescence periods, which were increased to 4-7 s for ITIs and 0.5-2 s for quiescence periods.

Training days were not always consecutive, and in instances where a day was skipped between training, mice were provided with a minimum amount of water in their home cage.

#### Passive stimulus presentation

Passive stimulus presentation was performed for 3 days before training to serve as acclimation to the recording rig and to provide a baseline response to visual stimuli. Passive stimuli were also presented after each session of task performance. The stimuli presented were gratings of the same size and spatial frequency as those presented during the task, but on either the left screen (never seen during the task), the center screen (seen during reward in the task), or the right screen (seen at the start of each trial during the task). The order of the three stimuli was randomized for each presentation, and all stimuli were presented 50 times. Stimuli were presented for 500 ms, with a 2-3 s inter-stimulus interval randomly chosen in 100 ms increments. All stimulus presentations with wheel movement were excluded from analysis.

#### Widefield imaging and fluorescence processing

Widefield imaging was conducted with a sCMOS camera (PCO Edge 5.5) affixed to a macroscope (Scimedia THT-FLSP) with a 1.0x condenser lens and 0.63x objective lens (Leica). Images were collected with Camware 4 (PCO) and binned in 2×2 blocks giving a spatial resolution of 20.6 μm/pixel at 70 Hz. Illumination was generated using a Cairn OptoLED with alternating colors, yielding a 35 Hz signal for each color. Blue light (470 nm, excitation filter ET470/40x) was used to capture GCaMP calcium-dependent fluorescence, and violet light (405 nm, excitation filter ET405/20x) was used to capture calcium-invariant hemodynamic occlusion. Excitation light was sent through the objective with a 3 mm core liquid light guide and dichroic mirror and emitted light was filtered (525/50-55) before the camera.

Widefield movies were compressed using singular value decomposition (SVD) of the form *F = USV^T^*. The input to the SVD algorithm *F* was the *pixels x time* matrix of fluorescence values, and the outputs were *U,* the *pixels x components* matrix of spatial components, *V*, the *time x components* matrix of temporal components, and *S,* the diagonal matrix of singular values. The top 2000 components were retained, and all orthogonally invariant operations (such as deconvolution and averaging) were carried out on the matrix *S * V* to save processing time and memory.

Hemodynamic effects on fluorescence were removed by subtracting a scaled violet illumination signal from the blue illumination signal. Fluorescence data was spatially downsampled 3-fold, filtered between 5-15 Hz to emphasize the heartbeat frequency, and sub-sample shifted to temporally align the alternating blue- and violet-illumination. A scaling factor was then regressed for each pixel from the violet to the blue illumination signal. The violet illumination signal was then multiplied by this scaling factor and subtracted from the blue illumination signal.

To correct for slow drift, hemodynamic-corrected fluorescence was then linearly detrended, high-pass filtered over 0.01 Hz, and ΔF/F_0_ normalized, where F_0_ was the average fluorescence by pixel across the session softened by the median fluorescence across pixels. Fluorescence was deconvolved using a kernel previously fit from simultaneous widefield imaging and electrophysiology (Peters et al., 2021).

To combine SVD-compressed widefield data across recordings, data was recast from experiment-specific SVD components into a master basis set of temporal components *U_master_*, which was previously created from the spatial components of many mice. After aligning the spatial components within an experiment *U_experiment_* to the master alignment, temporal components (*S * V*)_*experiment*_ for each experiment were recast by

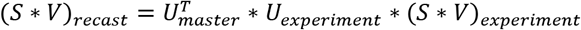

In this way, fluorescence data could be combined across experiments as temporal components *V* from a common basis set of spatial components *U_master_*, greatly reducing processing time and memory.

#### Retinotopic mapping

Cortical visual areas were mapped using visual sparse noise stimuli. White squares 7.5° in length were presented asynchronously on a black background, with each square lasting 166 ms and ~12% of squares being present at any given time. Activity for each square presentation was averaged within a 300-500 ms time window (corresponding to the maximum GCaMP6s signal), and the average response to each square was bootstrapped 10 times. Visual field sign maps (e.g. **Figure S2A**) were calculated for each bootstrapped mean and then averaged. Visual field sign was defined by gaussian-smoothing average square responses, finding the center-of-mass for each cortical pixel relative to the square azimuth and elevation locations, determining the gradient within these azimuth and elevation center-of-mass maps, and taking the sine of the difference between the azimuth and elevation gradients.

#### Widefield alignment

Widefield images were aligned across days within each mouse, and across mice to a master alignment, and to the Allen CCF atlas (Wang et al., 2020) (CCF v3, © 2015 Allen Institute for Brain Science) from the master alignment. Alignment across days was done using the average image within each day, first by finding vasculature edges through subtraction of a gaussian-blurred average image from the raw image, then by rigid-aligning these vasculature edges across days. Images were aligned across mice using retinotopic visual field sign maps. The visual field sign map for each mouse was affine aligned to a master visual field sign map previously created from an average and symmetrized map from many mice. The CCF atlas was aligned to the master visual field sign map by assigning expected visual sign values to each visual area on the atlas, then affine aligning the CCF visual sign map to the master visual sign map. Note the CCF alignment was only used to overlay area borders on widefield images and was not used for data processing.

#### Widefield movement-weighted hemisphere subtraction

We approximately isolated visually evoked fluorescence from movement-evoked fluorescence during task performance through a weighted hemisphere difference. Visually evoked activity is largely unilateral while movement evoked activity is largely bilateral (Shimaoka et al., 2019)(**Figure 2A**), which allows us to subtract movement evoked activity in the following manner. We make two assumptions, first, that stimulus and movement are additive, which previous well-fitting linear models support (Coen et al., 2021; Peters et al., 2021; Shimaoka et al., 2019; Steinmetz et al., 2019), and second, that the timecourse of visually and movement-evoked activity are the same in the left and right hemisphere, which is evident in our data. From this, the fluorescence timecourse *F*(*t*) in a given region can be described as a sum of unilaterally specific visual (*v*) and motor (*m*) gains for bilaterally symmetric visual- (*V*(*t*)) or motor- (*M*(*t*)) related activity through *F*(*t*) = *v*V*(*t*) + *m * M*(*t*). The fluorescence for a given region in the left and right hemisphere then will be

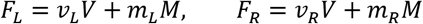

As mice often make movements during the delay period while no visual stimuli are present, we can estimate the ratio between left and right movement components *m_R_/m_L_*. This ratiometric approach assumes that, while the total fluorescence for each movement may change depending on factors like movement vigor, the ratio in fluorescence between the left and right hemispheres remains consistent during counterclockwise wheel movements. Using this ratio, we can then estimate a signal proportional to only the visual signal, without the movement signal, in the left hemisphere

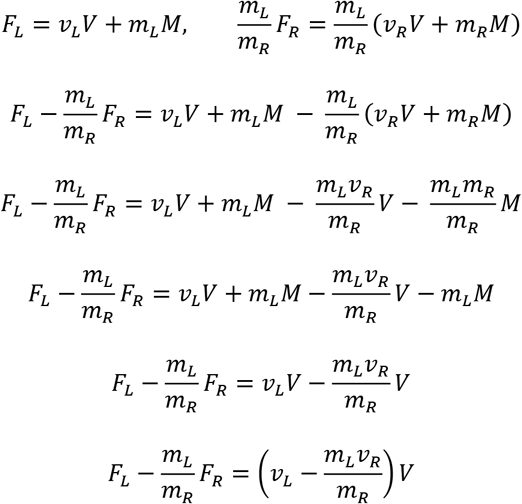

We determined the ratio *m_R_/m_L_* using movements during delay periods that would have been rewarded had a stimulus been present, i.e. reaching the reward threshold for leftward movement without reaching the punish threshold for a rightward movement. For each mouse, we averaged the fluorescence for these delay period movements within each day, then averaged the resulting fluorescence across days, then fit a scaling factor between movement-evoked activity in the right and left hemisphere.

#### Electrophysiological recordings

Electrophysiological recordings were performed using Neuropixels 3A probes affixed to custom rods and moved with micromanipulators (Sensapex). Some mice were recorded across multiple days, with the same insertion point being targeted across each day. Across 5 mice, one had 4 recordings, two had 2 recordings, and 2 had 1 recording. To determine the probe trajectory, probes were coated in dye on the first day of recording by dipping in DiI (ThermoFisher) 5-6 times with a few seconds of air drying between dips. Data was collected using Open Ephys (Siegle et al., 2017), spike-sorted using Kilosort 2 (Stringer et al., 2019), and units representing noise were manually removed using Phy (Rossant et al., 2016).

Probe trajectories were reconstructed from histology by aligning histological slices to the Allen CCF atlas and manually tracing the dye track using publicly available custom code (https://github.com/petersaj/AP_histology). The endpoint of the probe cannot be reliably determined through histology and there were likely slight variations of probe depth across days, so the depth of the probe for each recording was determined with electrophysiological markers. During each recording, the top recording sites of the probe were deliberately left outside of the brain to provide a demarcation for the cortical surface. This demarcation was then identified in the data using the LFP signal, where the brain-external channels were highly correlated with each other and not correlated with brain-internal channels. The surface of the cortex was defined where there was a large drop in cross-channel LFP correlation starting at the top of the probe. This often corresponded to the location where the first units were detected. By applying this recording-specific depth to the histology-aligned trajectory, we could then determine the brain area for each probe site.

Multiunit signals (**Figure 4C-D**) were obtained by combining all spikes within a given area in each recording. Single-unit analyses (**Figure 4E**) were performed on a subset of high-quality units determined using publicly available custom code (https://github.com/Julie-Fabre/bombcell).

#### Behavioral camera analysis

Eye camera analysis was done using DeepLabCut (Mathis et al., 2018) (https://github.com/DeepLabCut/DeepLabCut) with an available model trained on pupil videos (https://github.com/sylviaschroeder/PupilDetection_DLC). Four markers were used to track the pupil, with pupil diameter being the average length between each pair of opposing markers. Only time points with at least two pupil markers over 80% likelihood were used, while other time points (e.g. when there was a blink) were excluded.

Whisker analysis was done by aligning images of mouse faces across all experiments by control point registration, then defining a linear region-of-interest across the whiskers. The pixels in the whisker region-of-interest were extracted for each video frame, and whisker movement was defined as the absolute value of the difference between consecutive frames, summed across pixels.

### Quantification and statistical analysis

#### Stimulus response statistics

To test whether the mouse was reacting to the stimulus, it was necessary to test whether the mouse had a shorter reaction time to the stimulus compared to chance. Reaction time here is defined as the time between the visual stimulus onset and the next wheel movement. The analysis of this question is complicated by the fact that, even if wheel turns occurred at random times, an increased rate of random wheel turns would lead to an apparent decrease in median reaction times. We therefore require a method that ascertains whether reaction times are faster than would be expected if wheel turns occurred at times unrelated to the visual stimuli, while accounting for changes in total turn rates.

To do this, we used a method based on conditional randomization. Conditional randomization is a very simple statistical framework, that surprisingly has seen little use until recently (Candès et al., 2018; Hennessy et al., 2016). This approach can be used when we do not know the full probability distribution of the observed data *X*, but can specify a null hypothesis for its conditional distribution given a conditioning statistic *S*(*X*). We compare the value of a test statistic *T*(*X*) to a null distribution obtained by randomly sampling of *T*(*X*’) from this conditional distribution 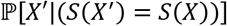, rejecting the null hypothesis if the actual test statistic exceeds the 95^th^ percentile of this distribution.

In our case, the full data *X* consists of the full wheel movement timeseries for all trials, together with each trial’s delay parameters *D_i_*: the lengths of the inter-trial-interval and quiescent period, which are randomly chosen for each trial. Knowing this data, we can compute the time *V_i_* when the visual stimulus appeared on each trial *i*, and the time *M_i_* of the next movement occurring after the visual stimulus on this trial. We define the conditioning statistic *S*(*X*) to be the wheel movement timeseries for all trials, together with the observed movement times *M_i_*. Thus, *S*(*X*) contains all information in *X* except for the delay parameters *D_i_*. We can thus sample from the conditional distribution 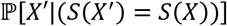 by rejection sampling: randomly redrawing the delay parameters for each trial, subject to the condition that the movement times that would have been detected with the new delay parameters are the same as those actually observed. For example, on a trial with a long delay between the stimulus onset *V_i_* and the next subsequent movement *M_i_*, the same movement time would have been registered if the ITI was longer and the stimulus later; but on a trial with a short delay between *V_i_* and *M_i_*, a longer ITI would have led to a stimulus coming later than *M_i_*.

We created a null set of reaction times by randomly resampling the delay parameters 10,000 times for each trial, subject to the condition that the observed movement time *M_i_* would still have been registered. Our test statistic *T*(*X*) was the median across trials of the reaction times *M_i_* – *V_i_*, excluding reaction times less than 100 ms as these fast reaction times were likely the result of coincidental timing rather than being related to the stimulus. Reaction times were considered significantly faster than chance at p < 0.05. The first day that reaction times were significantly faster than chance for each mouse was considered the first “association day”.

#### Single-unit response classification

Single units were classified as being significantly responsive to stimuli or movements through a shuffle test comparing firing rate in a baseline window with a response window (**Figure 4E**). For both stimulus and movement onset, baseline windows were set at 500-300 ms before each event, and response windows were set at 0-200ms after each event. Stimulus responses were obtained for all passive viewing trials with right-hand stimuli with no wheel movement. Movement responses were obtained from movements during task delay periods (with no stimulus present) that would have been rewarded had the stimulus been present (i.e., passing the counterclockwise threshold and not passing the clockwise threshold). The difference between baseline window and response window firing rate was compared to a null distribution, which was created by shuffling the baseline and response firing within each trial 1000 times, and cells were considered significantly responsive at p < 0.01.

### Data and code availability

The datasets generated during the current study are available as downloadable files at (FINAL DATA TO BE UPLOADED TO OSF BEFORE PUBLICATION).

The code used to analyze the data are available at (FINAL CODE TO BE UPLOADED TO GITHUB BEFORE PUBLICATION)

## Acknowledgements

We thank Charu Reddy and Bex Terry for animal husbandry, and Anne Ritoux and Yeqing Wang for histology. This work was supported by a Newton International Fellowship, EMBO Fellowship (ALTF 1428-2015), and a Human Frontier Science Program Fellowship (LL226/2016-L) to A.J.P., a Wellcome Trust PhD Studentship to J.M.J.F, and Wellcome Senior Investigator grants 205093 and 223144 to M.C and K.D.H. M.C. holds the GlaxoSmithKline/Fight for Sight Chair in Visual Neuroscience.

## Author contributions

A.J.P. conceived of and designed the study. A.J.P. and A.-M.M. collected and analyzed data, J.M.J.F. analyzed single-unit data. A.J.P., K.D.H., and M.C. wrote the manuscript with input from A.-M.M. and J.M.J.F.

---

**Figure S1.**
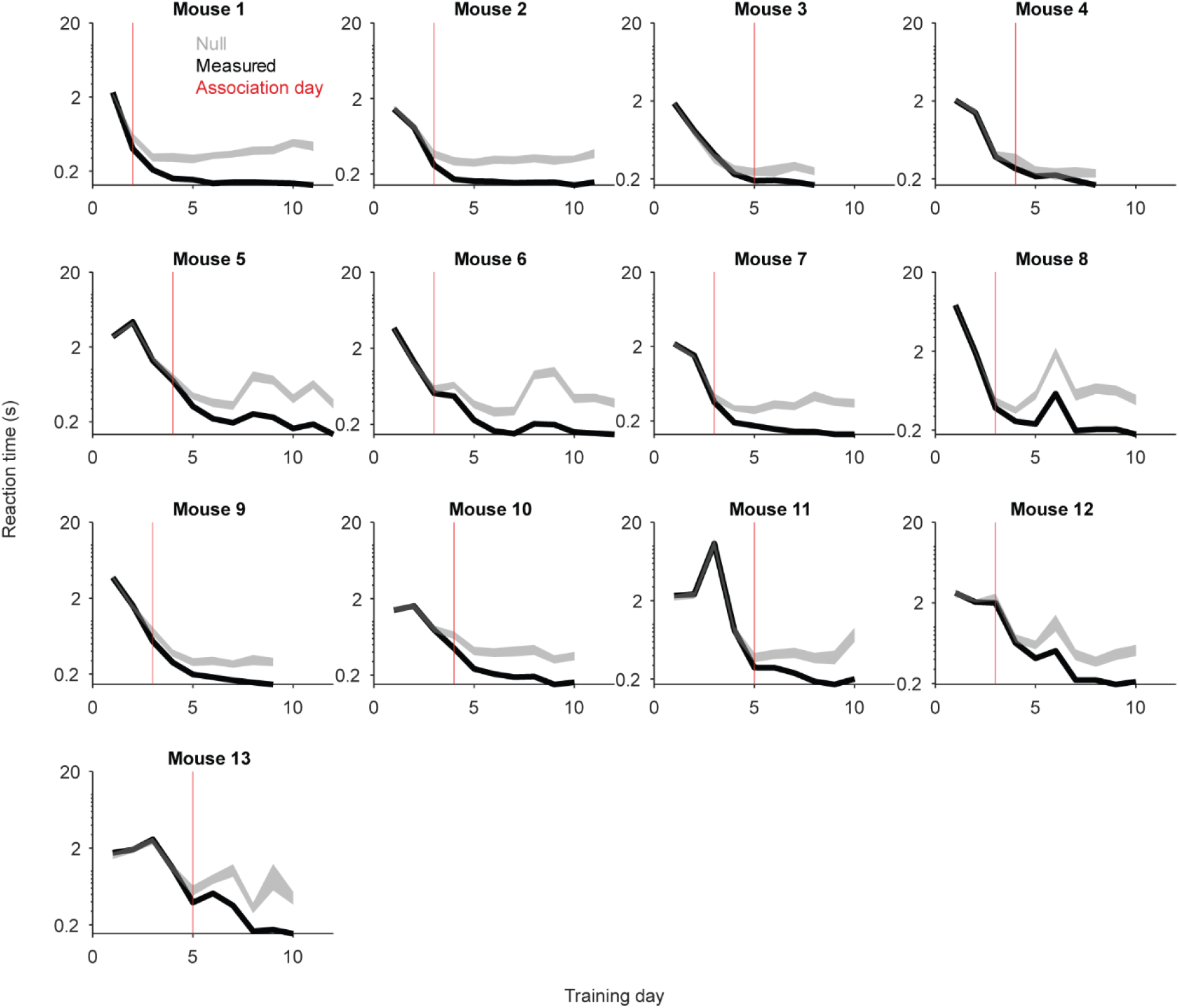
Individual animal reaction times. As in Figure 1F, median reaction times across days, measured (black) or expected from chance (gray) separately for all mice (n = 13 mice). Curves are measured values; shading shows 95% confidence intervals from the null distribution. Reaction times too fast to be stimulus-responsive (< 100ms) are excluded. Red lines indicate the first day that reaction times diverged from the null which represent the “association day”, which were used to split days by group in Figure 2 and to align by association day in Figure 3.

**Figure S2.**
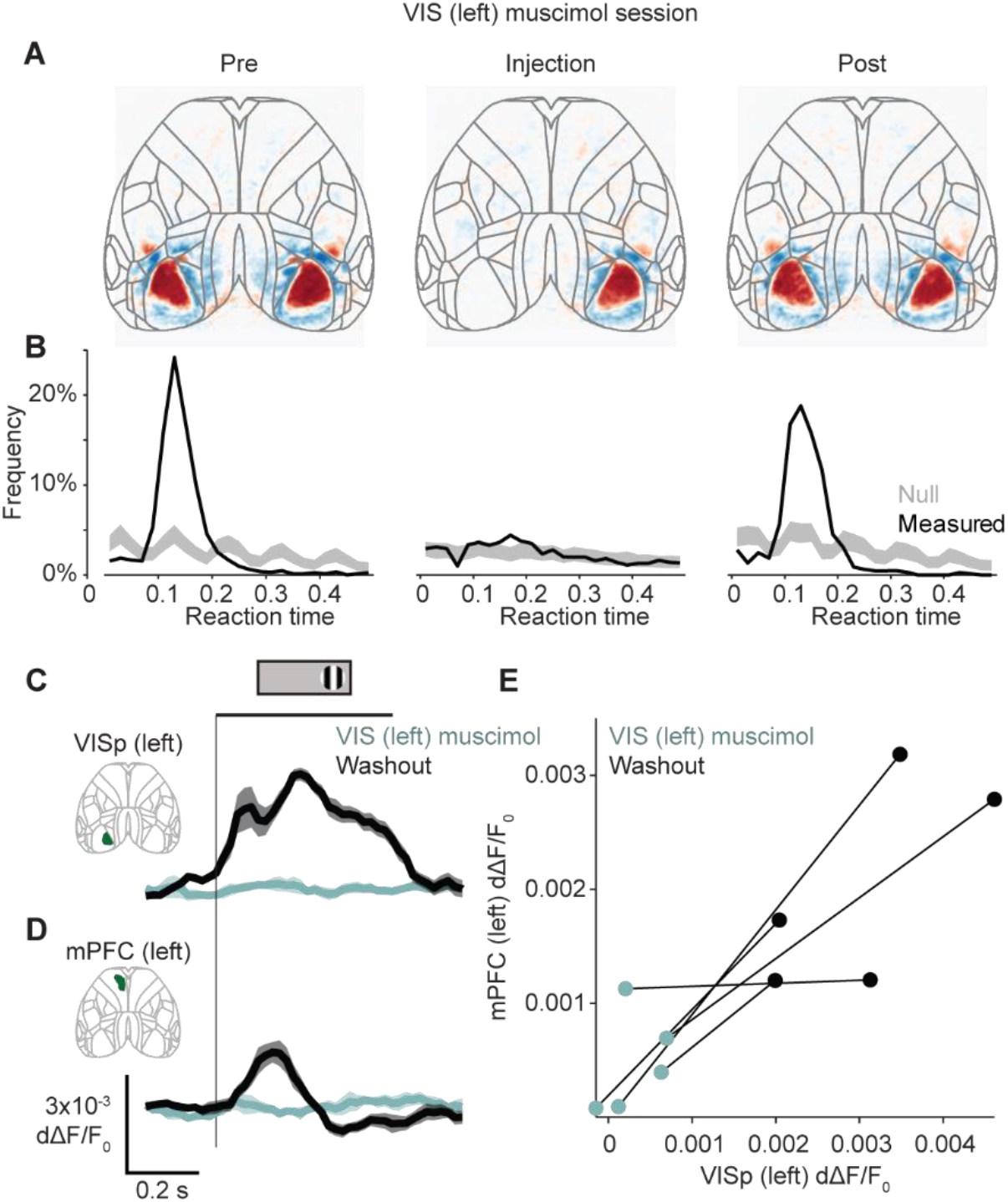
Muscimol abolishes visual and mPFC responses and movement timing without affecting total movement. (A) Visual field sign maps obtained from sparse noise visual presentation on days before (left), on (center), and after (right) muscimol injection in the left visual cortex averaged across mice (n = 5 mice). Muscimol injections into the left visual cortex abolishes retinotopic visual responses. (B) Histograms of reaction times measured (black) or expected from chance (null) as in Figure 1D. Curves are mean across mice and shadings are 95% confidence intervals from the null distribution (n = 5 mice). (C) Fluorescence in the left visual cortex during passive viewing of right-hand stimuli, on (cyan) and after (black) days with muscimol injection into the left visual cortex. Curves and error bars show mean ± s.e.m. across mice (n = 5 mice). (D) As in (C), for mPFC. (E) Fluorescence in the left visual cortex and mPFC as the maximum within 0-200 ms after stimulus onset, each connected pair is one mouse (n = 5 mice). Inactivating the visual cortex reduces stimulus-evoked responses in the mPFC (left-sided signed-rank test, p = 0.031).

**Figure S3.**
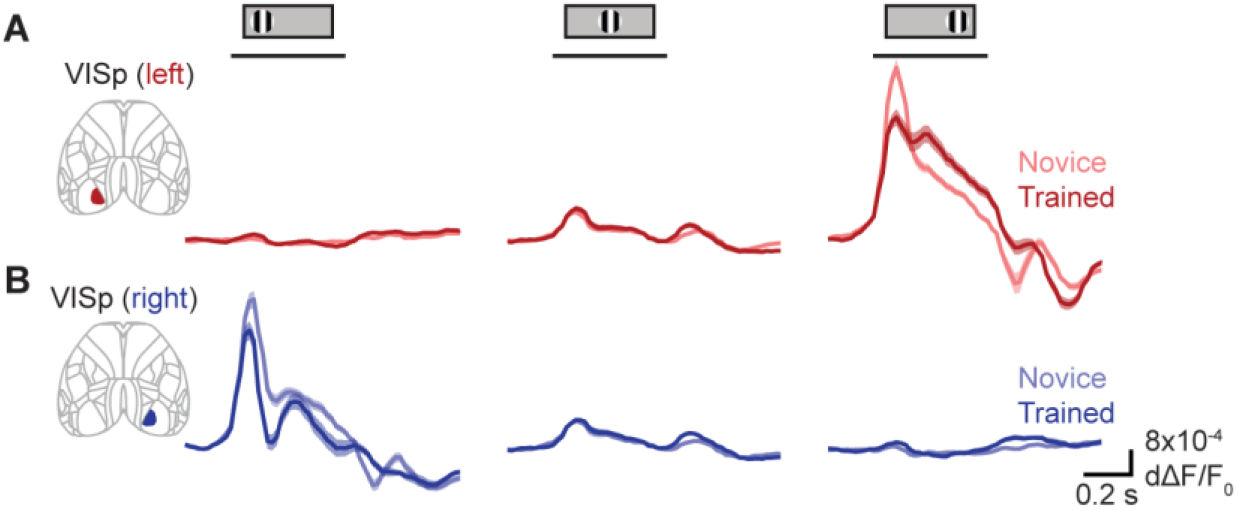
Left VISp fluorescence to right-hand stimulus becomes sustained with training. (A) Fluorescence during passive stimulus viewing in left hemisphere primary visual cortex (VISp) in novice (light red) and trained (dark red) mice. Curves and shading show mean ± s.e.m. across mice. Line under stimulus icon indicates when the stimulus is on the screen. Fluorescence in the left VISp to contralateral (right-hand) stimuli decreases in onset amplitude but becomes more sustained with learning (three-way ANOVA on time, learning stage, and stimulus, learning stage effect 1.2 x 10^-3^). (B) As in (A), for the right hemisphere VISp. Fluorescence in the right VISp to contralateral (left-hand) stimuli reduces in onset amplitude with learning (three-way ANOVA on time, learning stage, and stimulus, learning stage effect p = 0.013).

**Figure S4.**
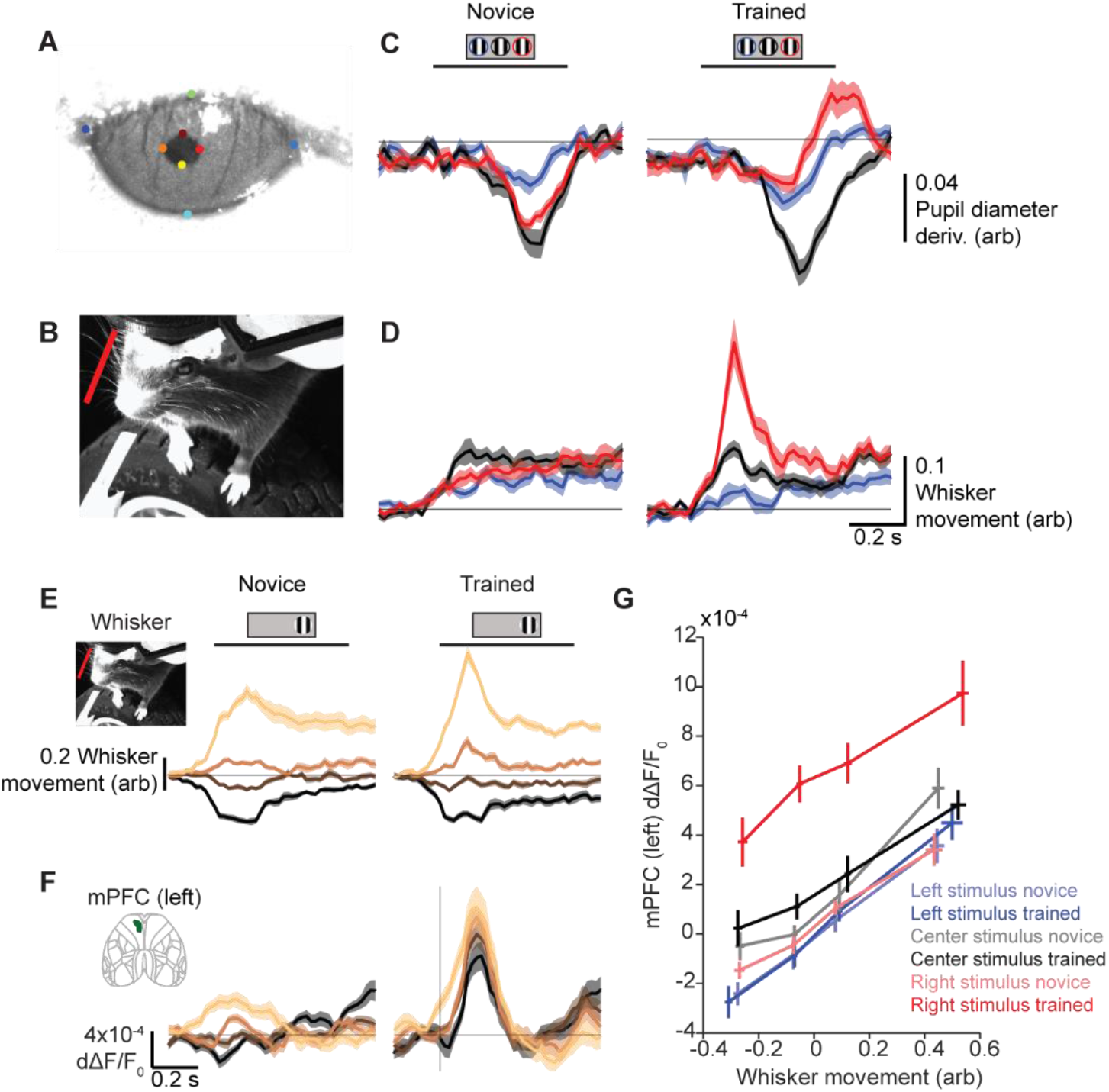
mPFC stimulus-evoked responses after learning are robust to behavioral changes. (A) Example eye video with points marked from DeepLabCut annotation. (B) Example face video with whisker region-of-interest in red. (C) Change in pupil diameter during passive viewing of stimuli on left (blue), center (black), and right (red) during novice (left) and trained (right) days. Curves are average ± s.e.m. across mice (n = 13). (D) As in (C), for whisker movement. (E) Whisker movement aligned to stimulus onset during novice (top) and trained (bottom) stages, binned into quartiles for each session by maximum movement 0-200 ms after stimulus onset. (F) Fluorescence in the left hemisphere mPFC from trials binned by whisker movement in (E). (G) Fluorescence in left mPFC by whisker movement, each averaged 0-200 ms after stimulus onset and binned by whisker movement in (E). Colors are stimuli on the left (blue), center (black), and right (red), for novice (pale) and trained (dark) stages. Curves are average ± s.e.m. across mice. Fluorescence increases in left mPFC specifically to right-hand stimuli regardless of movement (two-way ANOVA, stage effect left-hand stimulus p = 0.55, center stimulus p = 0.27, right-hand stimulus p = 1.3 x 10-^18^).

## REFERENCES

Allen, W.E., Kauvar, I. V., Chen, M.Z., Richman, E.B., Yang, S.J., Chan, K., Gradinaru, V., Deverman, B.E., Luo, L., and Deisseroth, K. (2017). Global Representations of Goal-Directed Behavior in Distinct Cell Types of Mouse Neocortex. Neuron 94, 891–907.e6.

Barthas, F., and Kwan, A.C. (2017). Secondary Motor Cortex: Where ‘Sensory’ Meets ‘Motor’ in the Rodent Frontal Cortex. Trends Neurosci. 40, 181–193.

Bhagat, J., Wells, M.J., Harris, K.D., Carandini, M., and Burgess, C.P. (2020). Rigbox: An open-source toolbox for probing neurons and behavior. ENeuro 7, 1–12.

Bichot, N.P., Schall, J.D., and Thompson, K.G. (1996). Visual feature selectivity in frontal eye fields induced by experience in mature macaques. Nature 381, 697–699.

Burgess, C.P., Lak, A., Steinmetz, N.A., Zatka-Haas, P., Bai Reddy, C., Jacobs, E.A.K., Linden, J.F., Paton, J.J., Ranson, A., Schröder, S., et al. (2017). High-Yield Methods for Accurate Two-Alternative Visual Psychophysics in Head-Fixed Mice. Cell Rep. 20, 2513–2524.

Candès, E., Fan, Y., Janson, L., and Lv, J. (2018). Panning for gold: ‘model-X’ knockoffs for high dimensional controlled variable selection. J. R. Stat. Soc. Ser. B Stat. Methodol. 80, 551–577.

Chen, T.W., Li, N., Daie, K., and Svoboda, K. (2017). A Map of Anticipatory Activity in Mouse Motor Cortex. Neuron 94, 866–879.e4.

Coen, P., Sit, T.P.H., Wells, M.J., Carandini, M., and Harris, K.D. (2021). Mouse frontal cortex mediates additive multisensory decisions. BioRxiv 34–42.

Costa, R., Cohen, D., and Nicolelis, M. (2004). Differential corticostriatal plasticity during fast and slow motor skill learning in mice. Curr. Biol. CB 14, 1124–1134.

Crochet, S., Lee, S.H., and Petersen, C.C.H. (2019). Neural Circuits for Goal-Directed Sensorimotor Transformations. Trends Neurosci. 42, 66–77.

Erlich, J.C., Bialek, M., and Brody, C.D. (2011). A cortical substrate for memory-guided orienting in the rat. Neuron 72, 330–343.

Ferezou, I., Haiss, F., Gentet, L.J., Aronoff, R., Weber, B., and Petersen, C.C.H.H. (2007). Spatiotemporal Dynamics of Cortical Sensorimotor Integration in Behaving Mice. Neuron 56, 907–923.

Glickfeld, L.L., Histed, M.H., and Maunsell, J.H.R. (2013). Mouse Primary Visual Cortex Is Used to Detect Both Orientation and Contrast Changes. 33, 19416–19422.

Goard, M.J., Pho, G.N., Woodson, J., and Sur, M. (2016). Distinct roles of visual, parietal, and frontal motor cortices in memory-guided sensorimotor decisions. Elife 5, 1–30.

Harris, J.A., Mihalas, S., Hirokawa, K.E., Whitesell, J. D., Choi, H., Bernard, A., Bohn, P., Caldejon, S., Casal, L., Cho, A., et al. (2019). Hierarchical organization of cortical and thalamic connectivity. Nature 575, 195–202.

Heilbronner, S.R., and Hayden, B.Y. (2016). Dorsal Anterior Cingulate Cortex: A Bottom-Up View. Annu. Rev. Neurosci. 39, 149–170.

Hennessy, J., Dasgupta, T., Miratrix, L., Pattanayak, C., and Sarkar, P. (2016). A Conditional Randomization Test to Account for Covariate Imbalance in Randomized Experiments. J. Causal Inference 4, 61–80.

Huda, R., Sipe, G.O., Breton-Provencher, V., Cruz, K. G., Pho, G.N., Adam, E., Gunter, L.M., Sullins, A., Wickersham, I.R., and Sur, M. (2020). Distinct prefrontal top-down circuits differentially modulate sensorimotor behavior. Nat. Commun. 11, 6007.

International Brain Laboratory, Aguillon-Rodriguez, V., Angelaki, D., Bayer, H., Bonacchi, N., Carandini, M., Cazettes, F., Chapuis, G., Churchland, A.K., Dan, Y., et al. (2021). Standardized and reproducible measurement of decision-making in mice. Elife 10, 1–28.

Kennerley, S., Behrens, T., and Wallis, J. (2011). Double dissociation of value computations in orbitofrontal and anterior cingulate neurons. Nat. Neurosci. 14, 1581–1589.

Komiyama, T., Sato, T.R., Oconnor, D.H., Zhang, Y.X., Huber, D., Hooks, B.M., Gabitto, M., and Svoboda, K. (2010). Learning-related fine-scale specificity imaged in motor cortex circuits of behaving mice. Nature 464, 1182–1186.

Lak, A., Okun, M., Moss, M.M., Gurnani, H., Farrell, K., Wells, M.J., Reddy, C.B., Kepecs, A., Harris, K.D., and Carandini, M. (2020). Dopaminergic and Prefrontal Basis of Learning from Sensory Confidence and Reward Value. Neuron 105, 700–711.e6.

Makino, H., Ren, C., Liu, H., Kim, A.N., Kondapaneni, N., Liu, X., Kuzum, D., and Komiyama, T. (2017). Transformation of Cortex-wide Emergent Properties during Motor Learning. Neuron 94, 880–890.e8.

Manita, S., Suzuki, T., Homma, C., Matsumoto, T., Odagawa, M., Yamada, K., Ota, K., Matsubara, C., Inutsuka, A., Sato, M., et al. (2015). A Top-Down Cortical Circuit for Accurate Sensory Perception. Neuron 86, 1304–1316.

Mathis, A., Mamidanna, P., Cury, K.M., Abe, T., Murthy, V.N., Mathis, M.W., and Bethge, M. (2018). DeepLabCut: markerless pose estimation of user-defined body parts with deep learning. Nat. Neurosci. 21.

Le Merre, P., Esmaeili, V., Charrière, E., Galan, K., Salin, P.-A., Petersen, C.C.H., and Crochet, S. (2018). Reward-Based Learning Drives Rapid Sensory Signals in Medial Prefrontal Cortex and Dorsal Hippocampus Necessary for Goal-Directed Behavior. Neuron 97, 83–91.e5.

Le Merre, P., Ährlund-Richter, S., and Carlén, M. (2021). The mouse prefrontal cortex: Unity in diversity. Neuron 109, 1925–1944.

Mohajerani, M.H., Chan, A.W., Mohsenvand, M., LeDue, J., Liu, R., McVea, D.A., Boyd, J.D., Wang, Y.T., Reimers, M., and Murphy, T.H. (2013). Spontaneous cortical activity alternates between motifs defined by regional axonal projections. Nat Neurosci 16, 1426–1435.

Moorman, D.E., and Aston-Jones, G. (2015). Prefrontal neurons encode context-based response execution and inhibition in reward seeking and extinction. Proc. Natl. Acad. Sci. U. S. A. 112, 9472–9477.

Mulder, A.B., Nordquist, R.E., Örgüt, O., and Pennartz, C.M.A. (2003). Learning-related changes in response patterns of prefrontal neurons during instrumental conditioning. Behav. Brain Res. 146, 77–88.

Murakami, T., Yoshida, T., Matsui, T., and Ohki, K. (2015). Wide-field Ca2+ imaging reveals visually evoked activity in the retrosplenial area. Front. Mol. Neurosci. 8, 1–12.

Murray, E.A., Bussey, T.J., and Wise, S.P. (2000). Role of prefrontal cortex in a network for arbitrary visuomotor mapping. Exp. Brain Res. 133, 114–129.

Orsolic, I., Rio, M., Mrsic-Flogel, T.D., and Znamenskiy, P. (2021). Mesoscale cortical dynamics reflect the interaction of sensory evidence and temporal expectation during perceptual decisionmaking. Neuron 1–15.

Otis, J.M., Namboodiri, V.M.K., Matan, A.M., Voets, E.S., Mohorn, E.P., Kosyk, O., McHenry, J.A., Robinson, J.E., Resendez, S.L., Rossi, M.A., et al. (2017). Prefrontal cortex output circuits guide reward seeking through divergent cue encoding. Nature 543, 103–107.

Peters, A.J., Fabre, J.M.J.J., Steinmetz, N.A., Harris, K.D., and Carandini, M. (2021). Striatal activity topographically reflects cortical activity. Nature 591, 420–425.

Peyrache, A., Khamassi, M., Benchenane, K., Wiener, S.I., and Battaglia, F.P. (2009). Replay of rule-learning related neural patterns in the prefrontal cortex during sleep. Nat. Neurosci. 12, 919–926.

Pinto, L., and Dan, Y. (2015). Cell-Type-Specific Activity in Prefrontal Cortex during Goal-Directed Behavior. Neuron 87, 437–450.

Reinert, S., Hübener, M., Bonhoeffer, T., and Goltstein, P.M. (2021). Mouse prefrontal cortex represents learned rules for categorization. Nature 593, 411–417.

Rossant, C., Kadir, S.N., Goodman, D.F.M., Schulman, J., Hunter, M.L.D., Saleem, A.B., Grosmark, A., Belluscio, M., Denfield, G.H., Ecker, A.S., et al. (2016). Spike sorting for large, dense electrode arrays. Nat. Neurosci. 19, 634–641.

Rushworth, M.F.S., Noonan, M.A.P., Boorman, E.D., Walton, M.E., and Behrens, T.E. (2011). Frontal Cortex and Reward-Guided Learning and Decision-Making. Neuron 70, 1054–1069.

Schneider, D.M., Sundararajan, J., and Mooney, R. (2018). A cortical filter that learns to suppress the acoustic consequences of movement. Nature 561, 1.

Shimaoka, D., Steinmetz, N.A., Harris, K.D., and Carandini, M. (2019). The impact of bilateral ongoing activity on evoked responses in mouse cortex. Elife 8, 1–19.

Siegle, J.H., López, A.C., Patel, Y.A., Abramov, K., Ohayon, S., Voigts, J., Black, C., Voigts, J., Buccino, A.P., Elle, M., et al. (2017). Open Ephys: An opensource, plugin-based platform for multichannel electrophysiology. J. Neural Eng. 14.

Siniscalchi, M.J., Phoumthipphavong, V., Ali, F., Lozano, M., and Kwan, A.C. (2016). Fast and slow transitions in frontal ensemble activity during flexible sensorimotor behavior. Nat. Neurosci.

Sreenivasan, V., Esmaeili, V., Kiritani, T., Galan, K., Crochet, S., and Petersen, C.C.H. (2016a). Movement Initiation Signals in Mouse Whisker Motor Cortex. Neuron 92, 1368–1382.

Sreenivasan, V., Kyriakatos, A., Mateo, C., Jaeger, D., and Petersen, C.C.H. (2016b). Parallel pathways from whisker and visual sensory cortices to distinct frontal regions of mouse neocortex. Neurophotonics 4, 1.

Steinmetz, N.A., Zatka-haas, P., Carandini, M., and Harris, K.D. (2019). Distributed coding of choice, action, and engagement across the mouse brain. Nature in press.

Stringer, C., Pachitariu, M., Steinmetz, N., Reddy, C.B., Carandini, M., and Harris, K.D. (2019). Spontaneous behaviors drive multidimensional, brainwide activity. Science 364, 255.

Sul, J.H., Jo, S., Lee, D., and Jung, M.W. (2011). Role of rodent secondary motor cortex in value-based action selection. Nat. Neurosci. 14, 1202–1208.

Svoboda, K., and Li, N. (2018). Neural mechanisms of movement planning: motor cortex and beyond. Curr. Opin. Neurobiol. 49, 33–41.

Tennant, K.A., Adkins, D.L., Donlan, N.A., Asay, A.L., Thomas, N., Kleim, J.A., and Jones, T.A. (2011). The organization of the forelimb representation of the C57BL/6 mouse motor cortex as defined by intracortical microstimulation and cytoarchitecture. Cereb. Cortex 21, 865–876.

Wal, A., Klein, F.J., Born, G., Busse, L., and Katzner, S. (2021). Evaluating visual cues modulates their representation in mouse visual and cingulate cortex. J. Neurosci. 41, 3531–3544.

Wang, Q., Ding, S.L., Li, Y., Royall, J., Feng, D., Lesnar, P., Graddis, N., Naeemi, M., Facer, B., Ho, A., et al. (2020). The Allen Mouse Brain Common Coordinate Framework: A 3D Reference Atlas. Cell 181, 936–953.e20.

Wekselblatt, J.B., Flister, E.D., Piscopo, D.M., and Niell, C.M. (2016). Large-scale imaging of cortical dynamics during sensory perception and behavior. J. Neurophysiol. 115, 2852–2866.

Yang, G., Lai, C.S.W., Cichon, J., Ma, L., Li, W., and Gan, W.-B. (2014). Sleep promotes branch-specific formation of dendritic spines after learning. Science (80-.). 344, 1173–1178.

Zatka-Haas, P., Steinmetz, N.A., Carandini, M., and Harris, K.D. (2021). Sensory coding and the causal impact of mouse cortex in a visual decision. Elife 10, 501627.

Zhang, S., Xu, M., Kamigaki, T., Hoang Do, J.P., Chang, W.-C., Jenvay, S., Miyamichi, K., Luo, L., and Dan, Y. (2014). Long-range and local circuits for top-down modulation of visual cortex processing. Science (80-.). 345, 660–665.

